# The impact of brain lesions on tDCS-induced electric field magnitude

**DOI:** 10.1101/2021.03.19.436124

**Authors:** Ainslie Johnstone, Catharina Zich, Carys Evans, Jenny Lee, Nick Ward, Sven Bestmann

## Abstract

**Background:** Transcranial direct current stimulation (tDCS) has been used to enhance motor and language rehabilitation following a stroke. However, improving the effectiveness of clinical tDCS protocols depends on understanding how lesions may influence tDCS-induced current flow through the brain.

**Objective:** We systematically investigated the effect of brain lesions on the magnitude of electric fields (e-mag) induced by tDCS, and how to overcome lesion-induced inter-individual variability in e-mag.

**Methods:** We simulated the effect of 630 different lesions - by varying lesion location, distance from the target region of interest (ROI), size and conductivity - on tDCS-induced e-mag in the brains of two participants. Current flow modelling was conducted for two tDCS montages commonly used in clinical applications, which target either primary motor cortex (M1) or Broca’s area (BA44), respectively. We further explored how the inherent variability in e-mag that is introduced by inter-lesion differences can be overcome by individualising tDCS protocols.

**Results:** The effect on *absolute* e-mag was highly dependent on lesion size, conductance and the distance from the target ROI. Larger lesions, with high conductivity, closer to the ROI caused e-mag changes of more than 30%. The *sign* of this change was determined by the location of the lesion. Specifically, lesions located in-line with the predominant direction of current flow increased e-mag in the ROI, whereas lesions located in the opposite direction caused a decrease. Lesions had a large impact on the optimal electrode configuration if attempting to maximise for the total e-mag in the ROI, but little impact if only the component of e-mag flowing radially inward to the cortex was maximised. Knowing the effect of a given lesion on e-mag also allows for individualising tDCS intensity to reduce variability.

**Conclusions:** These results demonstrate that tDCS-induced electric fields are profoundly influenced by lesion characteristics, and further exacerbate the known variability in e-mag across individuals. Additionally, the dependence of these results on the assigned conductance of the lesion underlines the need for improved estimates of lesion conductivity for current flow models. Our results highlight the need for individualised dose control of tDCS in the lesioned brain to overcome the substantial inter-individual variability in electric fields delivered to a cortical target region.

**Highlights:**

- Lesions can alter tDCS-induced electric field magnitude (e-mag) in a target by 30%
- Lesions can cause increases or decreases to e-mag
- Direction of change depends on the position of the lesion relative to current flow
- Lesion conductivity - the true value for which is unknown - also impacts change
- E-mag variability can be reduced by individualising montage and stimulation intensity

## Introduction

Transcranial direct current stimulation (tDCS) is proposed as an economical and non-invasive method of enhancing recovery after stroke when paired with behavioural training (Allman et al., 2016; Butler et al., 2013; Galletta et al., 2016; Holland et al., 2011; Holland and Crinion, 2012; Hummel et al., 2005, 2006; Johnstone et al., 2017; Kang et al., 2015; Stagg et al., 2012). However, the effects of tDCS vary substantially across individuals (Elsner et al., 2016; Wiethoff et al., 2014) making adoption into routine clinical practice difficult. Individual differences in brain and skull anatomy lead to large variations in how much current reaches the target brain regions (Evans et al., 2020; Laakso et al., 2015). This leads to unacceptable variability in the physiological and behavioural effects of tDCS across individuals (Elsner et al., 2016; Wiethoff et al., 2014). Variability is likely to be further exacerbated in stroke patients, who constitute end-users of tDCS, where lesions may distort current flow (Minjoli et al., 2017). For example, it is highly likely that lesion size and location could influence current flow and are unlikely to be identical in any two patients. Moreover, while it is known that lesions have different conductance compared to healthy brain tissue (McCann et al., 2019) the magnitude of this difference is not known, and it remains unclear how this affects current flow.

The amount of current, and the path it takes through an individual’s brain, can be estimated using current flow modelling (Huang et al., 2017; Thielscher et al., 2015). Here, high-resolution volume conduction models are usually based on individual MRI scans, to account for the complex geometry of the head and brain. From these MR images, the head is segmented into different tissue types, each with an assigned conductivity, and the volumetric anatomical images are tessellated into a 3D mesh. The voltage distribution for the resulting finite element model (FEM) is then obtained by numerically solving the Laplace equation (Datta et al., 2009). Current flow models can be used to estimate variability across individuals (Laakso et al., 2015), determine the individual stimulation intensities required to generate equivalent electric fields across participants (Evans et al., 2020), or even optimise the electrode location to target specific regions (Dmochowski et al., 2013, 2011; Huang et al., 2018).

These models have been applied to the brains of stroke patients (Datta et al., 2011; Dmochowski et al., 2013; Galletta et al., 2015; Minjoli et al., 2017; Piastra et al., 2021) with some studies finding profound effects of the lesioned tissue on current flow (Datta et al., 2011; Minjoli et al., 2017; Piastra et al., 2021). Despite these efforts, the multi-faceted nature of lesion characteristics and the complexities of their influence(s) on current flow remain largely unknown. Previous studies are based on the results from a small number of example lesions. However, while lesions tend to be confined to certain vascular territories across patients, there is a large variation in lesion location within these regions, and in principle, all areas of the brain have the potential to be affected by a stroke (Kim et al., 2019). In addition to variability in lesion location, it is known that the size of lesions varies dramatically. However, despite large variations in lesion size patients may have very similar symptoms (Hope et al., 2013; Zhao et al., 2018), meaning study or treatment groups are unlikely to have homogenous lesions. This makes it difficult to generalise the conclusions of these previous modelling studies and make decisions on how to incorporate these findings into protocols for individual patients with potentially very different lesions.

A further issue is that the validity of results from current flow modelling simulations rely on the accuracy of tissue segmentation and choice of correct conductance values. Previous studies have opted to model lesions with conductance equivalent to cerebral spinal fluid (CSF) (Datta et al., 2011; Dmochowski et al., 2013; Galletta et al., 2015; Minjoli et al., 2017) but the actual conductance value of lesioned tissue is not known. Estimates obtained from various MRI techniques vary 10-fold (McCann et al., 2019), ranging from values below that of typical grey and white matter conductance, to above the value typically assigned to CSF. The influence that changes in lesion conductance have on current flow simulations remains unclear.

In this study, we systematically assessed the influence of synthetic lesions on induced electric field magnitude within two regions of interest (ROIs): M1 and BA44 – two regions commonly targeted by tDCS for clinical applications. We used synthetic lesions to systematically vary the location of the lesion, its distance to an ROI, its size, and its conductance, to determine general patterns or rules that govern how lesions might alter current flow. We further investigated approaches to compensate for the lesion-induced variability in electric field magnitude. Specifically, we explored the effect of individualising tDCS montages and stimulator intensity to optimise electric field magnitude in the ROI, and thereby reduce variability in e-mag between cohorts with either lesioned or healthy brains. Together, our approach and findings can be used to guide the application of tDCS for diverse study populations and emphasises the need for individualized application of tDCS in populations affected by brain damage.

## Methods

### Overview

A variety of synthetic spherical lesions were modelled in the brains of two example participants, for two different electrode montages with utility in stroke rehabilitation research. For each montage an ROI was defined-motor cortex (M1) and Broca’s area (BA44)- and lesions differed in their cardinal direction from the ROI, distance from the ROI and the lesion’s size. For each lesion, simulations were run using a range of conductance values.

### Participants

The T1-weighted 3D structural MR scans of two healthy females were used in this analysis, both of whom gave informed consent to have their scans used for this purpose. The project was approved by the UCL Research Ethics Committee (project no: 14233/001). P01 was 26 years old at the time of scanning, P02 was 25 years old. Both scans took place at the Wellcome Centre for Human Neuroimaging at UCL (WCHN, UCL) on a Siemens 3 Tesla TIM Trio scanner with a 64-channel head coil (176 sagittal slices, matrix size 256 x 256, 1mm isotropic resolution, TR/TE=1900/3.96).

### Current flow modelling

Current flow modelling was performed using a custom version of ROAST 2.7 (Huang et al., 2019). ROAST is a fully automated, open-source MATLAB application that performs segmentation of the brain image (via SPM), places virtual stimulation electrodes, performs volumetric meshing (via Iso2Mesh-Fang & Boas, 2009) and solves the finite element model numerically to estimate current flow. By default, ROAST segments the image into 6 tissue types (skin, bone, CSF, grey matter, white matter and air). We modified ROAST (i.e. ROAST-lesion) to incorporate a seventh tissue type (lesion). Consequently, volumetric meshing and solving the finite element model were performed using the 7-tissue head model. With the exception of the additional lesion and the tDCS montage information ROAST 2.7 default settings were used for all simulations. Code for ROAST-lesion is available at [*code provided upon acceptance*].

### tDCS stimulation montages

Two montages with utility in stroke rehabilitation research were selected for this study. One typically used in motor rehabilitation research (Amadi et al., 2015; O’Shea et al., 2014) to target M1, where the anode is applied over left primary motor cortex (M1) and the cathode over right supraorbital ridge. Specifically, we placed electrodes over 10-05 coordinates Fp2 and CCP3 for both participants. For the other montage, often used in aphasia rehabilitation studies (Galletta et al., 2016; Holland and Crinion, 2012) to target BA44, the anode is positioned over left frontal cortex (BA44) and the cathode on the right neck. Specifically, we placed the electrodes over Exx20 and FFT7h for participant one (P01) and over Exx20 and F7h for participant two (P02).

The choice of locations used here are intended to be taken as examples of potential montages. In reality, for some tDCS studies the exact locations of stimulation are individualised either to target intact tissue or to target anatomically or functionally defined regions (Baker et al., 2010; Fridriksson et al., 2011). All simulations were run with an applied current strength of 1mA. It should be noted that, in line with Ohm’s Law, e-mag values will scale linearly with increases in applied current given constant conductance. All simulations used disk electrodes of radius 17mm and depth 2mm.

### Lesions

Synthetic lesions were positioned relative to the ROIs. ROIs were 12mm radius spheres centred at manually defined M1 hand knob or BA44. All lesions were spherical and constrained to the grey and white matter. The lesions were positioned relative to their cardinal direction from the ROI; either due right (R), left (L), anterior (A), posterior (P), superior (S), inferior (I), right-anterior-superior (RAS), right-anterior-inferior (RAI), right-posterior-superior (RPS), right-posterior-inferior (RPI), left-anterior-superior (LAS), left-anterior-inferior (LAI), left-posterior-superior (LPS), or left-posterior-inferior (LPI) of the ROI (see Figure 1). Lesions varied in distance from the ROIs (shortest Euclidean distance from the edge of ROI to the edge of lesion: 1mm, 5mm, 10mm), and in size (radii: 4mm, 12mm, 24mm). Lesions were not designed to be morphologically realistic, but rather to ensure comparability and quantification of the impact on electrical fields.

**Figure 1:**
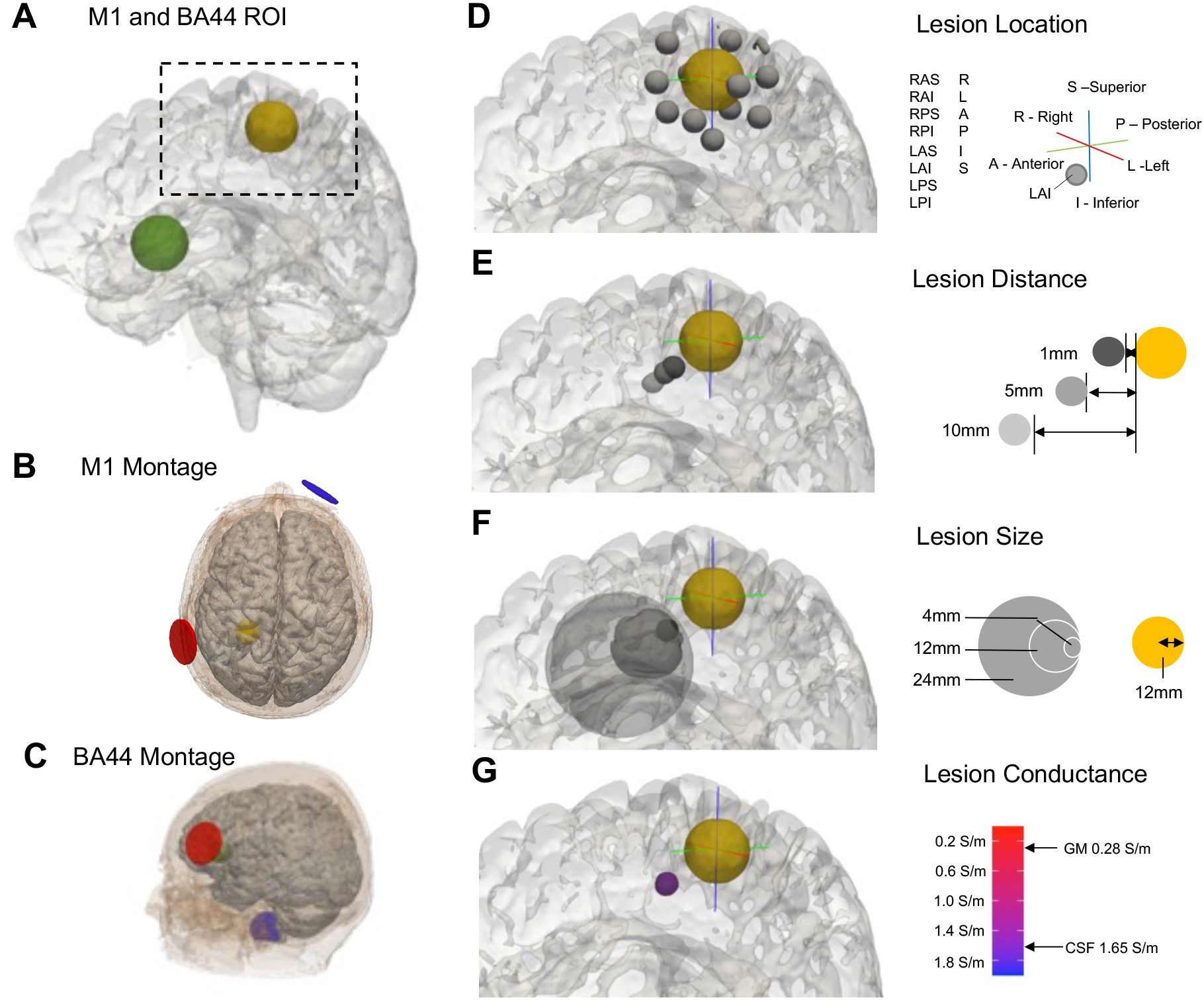
Modelled lesion parameters. A: Rendered brain of example participant (P01) with M1 ROI (yellow) and BA44 ROI (green). B: tDCS montage targeting M1, with anode in red, cathode in blue. C: BA44 tDCS montage. D: The 14 potential lesion directions N.B. lesions where the central voxel was outside the brain were omitted. E: The three different lesion distances (1mm, 5mm, 10mm), where distance was measured as the shortest Euclidean distance from the edge of the ROI to the edge of the lesion. F: The three different lesion sizes, defined by lesion radius (4mm, 12mm, 24mm). G: The five different lesion conductivities (0.2S/m, 0.6S/m, 1S/m, 1.4S/m, 1.8S/m)

Current flow simulations were only performed if the centre of the lesion was within the brain, and if the volume of the lesion was at least 20% of the maximum potential volume. If these conditions were met, then simulations were run with a variety of different lesion conductivities (0.2 S/m, 0.6 S/m, 1 S/m, 1.4 S/m, 1.8 S/m) which ranged from roughly the conductance of grey and white matter (0.28 S/m and 0.13 S/m respectively) to above that of CSF (1.65 S/m), spanning the range of values reported in McCann et al., 2019. Out of a possible 630 simulations, a total of 490 and 409 M1 simulations, and 435 and 355 BA44 were run for P01 and P02 respectively. Note that there are fewer simulations for the BA44 montage due to the close proximity of the lesion to the cortical surface.

### Simulation outputs and independent variables

3D images of electric field magnitude (e-mag) values, measured in V/m, were extracted for each lesion simulation. Additionally, for each participant, and both montages, a simulation without lesions was performed to obtain estimates of e-mag within the ‘healthy’ non-lesioned brain. The e-mag change for each lesion was calculated by subtracting the non-lesioned e-mag image from the lesioned e-mag image. The mean, the 16^th^ percentile and the 84^th^ percentile e-mag change were computed across the grey matter voxels within the target ROI (see Figure 2).

Furthermore, to calculate the direction of the electric field relative to the ROI, the component of the electric field magnitude in the x, y, and z plane was extracted from the non-lesioned simulation, for each participant in both montages. The electric field direction vector was calculated, for each participant in both montages, using the mean electric field in each plane from all grey matter voxels within the target ROI.

Lesion distance, size and conductivity were treated as continuous numerical variables in all analyses. In order to quantify lesion location and allow for comparison between montages, the angle between i) the 3D electric field direction vector from within the grey matter of the ROI in the non-lesioned simulation (discussed in paragraph above), and ii) the 3D direction vector indicating the ‘movement’ from the ROI to the lesion, was calculated (see Figure 2E). This angle indicates the degree to which the lesion was located in the path of current flow, e.g., a small angle indicates the lesion is in the path of current flow.

### Identifying optimal montages

To assess whether altering the tDCS montage could compensate for lesion-induced variability in electric field magnitude we determined the optimal tDCS montage for selected lesions using the M1 montage on one participant (P01). To this end, we used the roast-target function of ROAST v3.0 (Dmochowski et al., 2013, 2011; Huang et al., 2018). Specifically, large (24mm radius), close (1mm distance) lesions located right-anterior-inferior (RAI) and left-posterior-inferior (LPI) were compared with the non-lesioned brain. These lesions were treated as having conductance equal to that of CSF (1.65 S/m), the stimulator output current was set at 1mA and optimisation was constrained to a bipolar montage. For each condition roast-target optimisation was run twice with two different aims. First, the objective function was to maximise the total e-mag within the grey matter of the M1 ROI. Second, we maximised specifically the component of the e-mag flowing radially inward through the cortex. Both options have been discussed as viable choices for optimization of tDCS with current flow modelling (Dmochowski et al., 2011).

### Analysis

All MRI manipulations were performed using tools from the FMRIB software library (FSL; Jenkinson et al., 2012) and all data analysis was performed using R v4.0.3 in RStudio v1.3.1093.

## Results

### Larger lesions, closer to the ROI, with higher conductivity have a greater impact on e-mag change within ROIs

First, we evaluated the effects of lesion distance, size and conductivity on the *absolute* mean e-mag change of the ROI. To this end we used two linear mixed effect models, one for the M1 montage and one for the BA44 montage, with participant as a random effect impacting intercept. For both montages, lesions with closer distance (M1: F(1,894)=16.9, β=−0.99, p=4.3e-5, Figure 3A; BA44: F(1,785)=46.6, β=−1.05, p=2.1e-11, Figure 3D), larger size (M1: F(1,894)=90.4, β=2.31, p<2.2e-16, Figure 3B; BA44: F(1,785)=178, β=2.11, p<2.2e-16, Figure 3E), and higher conductance (M1: F(1,894)=27.4, β=2.25, p=1.4e-10, Figure 3C; BA44: F(1,785)=54.6, β=1.63, p=3.8e-13, Figure 3F) had a greater impact on absolute change in e-mag within the ROI.

**Figure 3:**
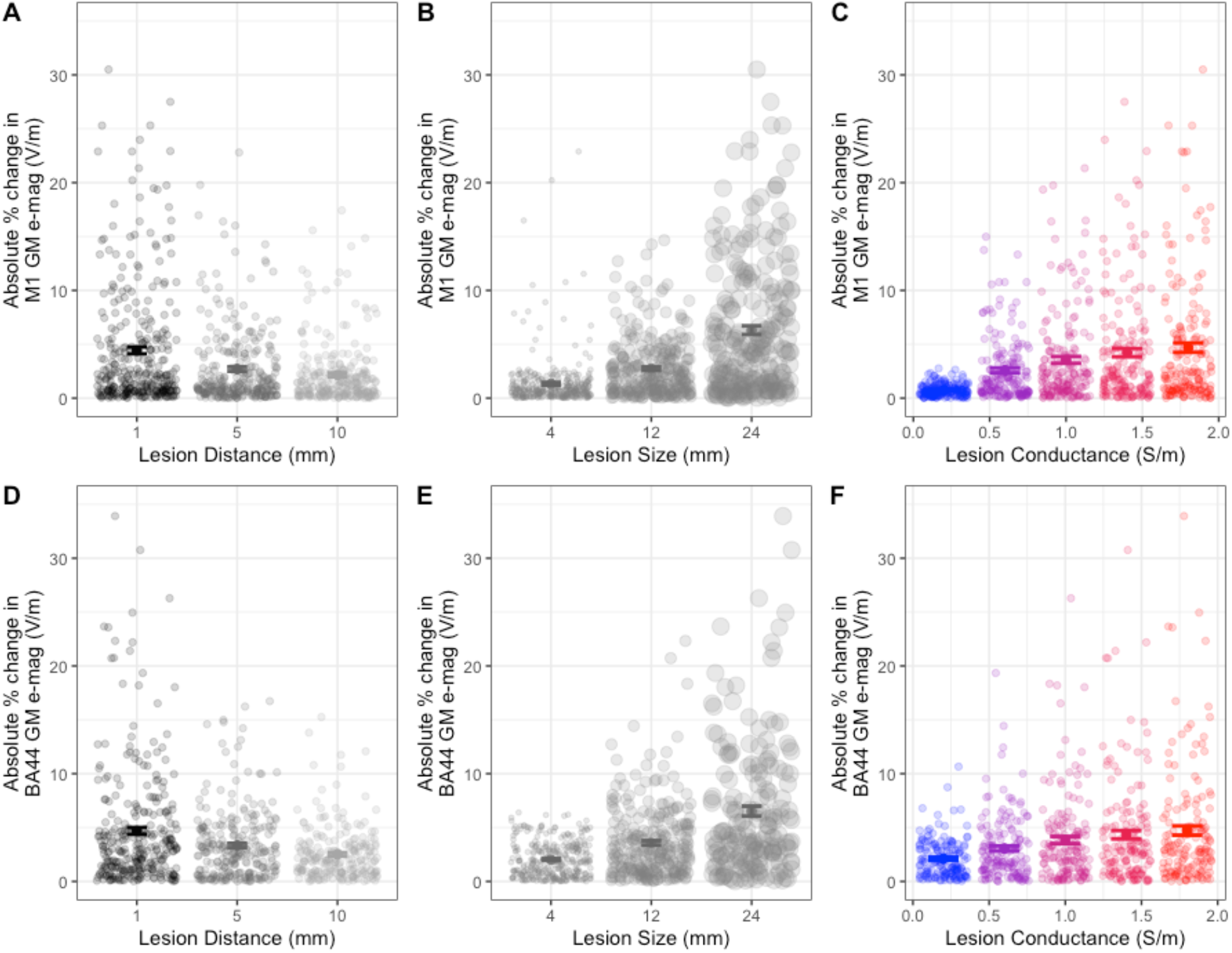
Larger lesions, closer to the ROI, with high conductivity have a greater impact on electric field magnitude (e-mag) change within the ROI. A-C: Scatter plots with mean and standard error (SE) of the *absolute* percentage e-mag changes in M1 caused by lesions with different sizes, distances and conductance. Data are the results from individual simulations and pooled from both participants for analysis. D-F: Scatter plots with mean and SE of *absolute* e-mag change in BA44 GM. Individual data points are jittered on the x-axis for display purposes.

There was no change in the significance of results if data from both montages/ROIs were included together in one linear mixed model with montage included as an additional random effect. Adding montage as a fixed effect also did not change results, and there were no significant interactions between montage and the other variables (p>0.14). This finding indicates that the effect of lesion distance, size or conductivity on e-mag change were comparable for the M1 and BA44 montage/ROI.

### Lesions in the path of current flow increase e-mag, while those in the opposite direction cause e-mag decreases

We went on to investigate the main effect of lesion location on the mean e-mag change in the ROIs, this time accounting for *sign*. Lesions due R, RAS and RAI of left M1 caused increases in M1 e-mag, whereas lesions due inferior, LPI, LAI or RPI tended to decrease M1 e-mag (see Figure 4 for selected examples, and Supplementary Figure 1 for data from all M1 simulations). For the BA44 montage however, lesions due R and RPI tended to increase e-mag within the ROI, whereas lesions due posterior and RAS tended to decrease e-mag (see Supplementary Figure 2 for data from all BA44 simulations).

**Figure 4:**
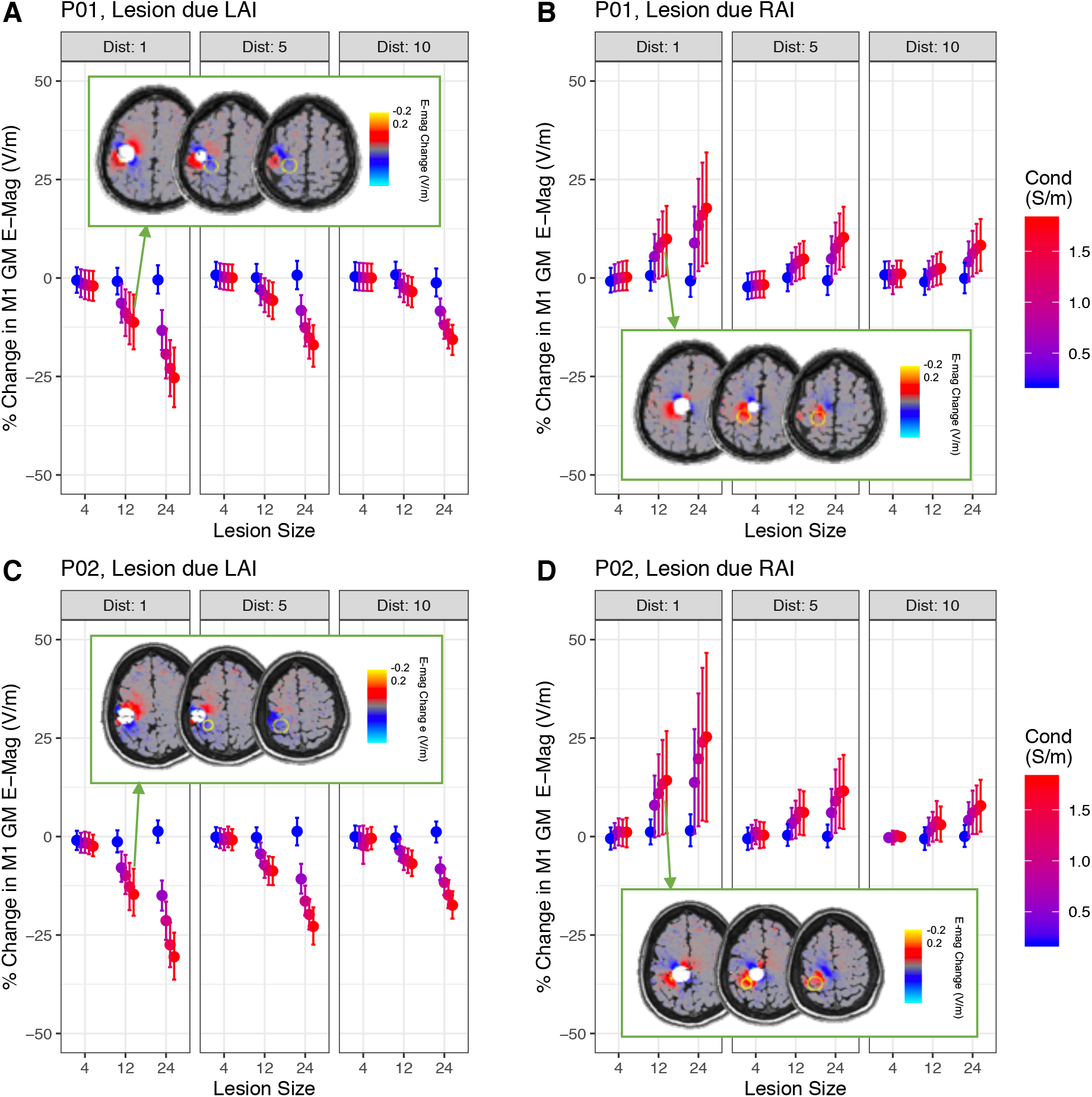
Lesion location determined the sign of electric field magnitude (e-mag) change. A, B: Percentage change in e-mag in M1 region of interest (ROI) caused by lesions due left-anterior-inferior (LAI) or right-anterior-inferior (RAI), with varying distances, sizes and conductivities for participant P01. Data points show mean percentage change across all ROI voxels, error bars show the 16^th^ and 84^th^ percentile values for change across voxels. Inset: Absolute change in e-mag across the whole brain for LAI and RAI lesions of size 12mm, distance 1mm and conductance 1.4S/m. C,D: Same as above, but for participant P02.

In order to quantify lesion location, and allow comparison between the two montages, the angle of lesion direction relative to the current flow in the ROI was calculated. This was achieved by obtaining the angle between i) the 3D vector describing the electric field direction within the ROI in the non-lesioned, and ii) brain the 3D vector linking the centre of the ROI with the centre of the lesion. Angle of lesion location was a significant predictor of percentage change in e-mag in the M1, as assessed by linear mixed model (t(897)=−9.31, β=−0.06, <2.2e-16), with participant as a random effect on intercept. The same relationship was found for the BA44 montage (t(788)=−13.3, β=−0.07, p<2.2e-16). This indicates that lesions that are more aligned with the direction of current flow in the ROI tended to increase mean e-mag in the ROI, whereas those that were located in the opposite direction to current flow tended to decrease e-mag.

Combining data from both montages, with montage as a random effect on intercept, did not change the significance of the effect of lesion location. Including montage as a fixed effect also did not result in a significant main effect of montage, or significant interaction between montage and lesion angle (p>0.13), again indicating that the results are consistent across the M1 and BA44 simulations.

### Interactions with lesion location

In order to assess whether the effect of lesion location was modulated by the other lesion characteristics, we performed additional linear mixed models. These included fixed effects of angle of lesion location relative to current flow direction in the ROI, distance, size and conductivity, as well as interactions between the angle of lesion location and the other variables. Participant was again included as a random effect on intercepts.

In conformity with the results outlined above, all main effects were significant. Further significant interactions between lesion location and distance (M1: t(891)=5.27, β=0.04, p=1.7e-7; BA44: t(782)=6.83, β=0.04, p=1.7e-11), lesion location and size (M1: t(891)=−12.7, β=−0.09, p<2.2e-16; BA44: t(782)=−13.0, β=−0.11, p<2.2e-16), and lesion direction and conductance (M1: t(891)=−6.02, β=−0.06, p=2.6e-9; BA44: t(782)=−9.53, β=−0.08, p<2.2e-16) were however found. This indicates that lesion direction had a greater effect for lesions with larger size, smaller distance from the ROI and higher conductance (see Figures 5 & 6).

**Figure 5:**
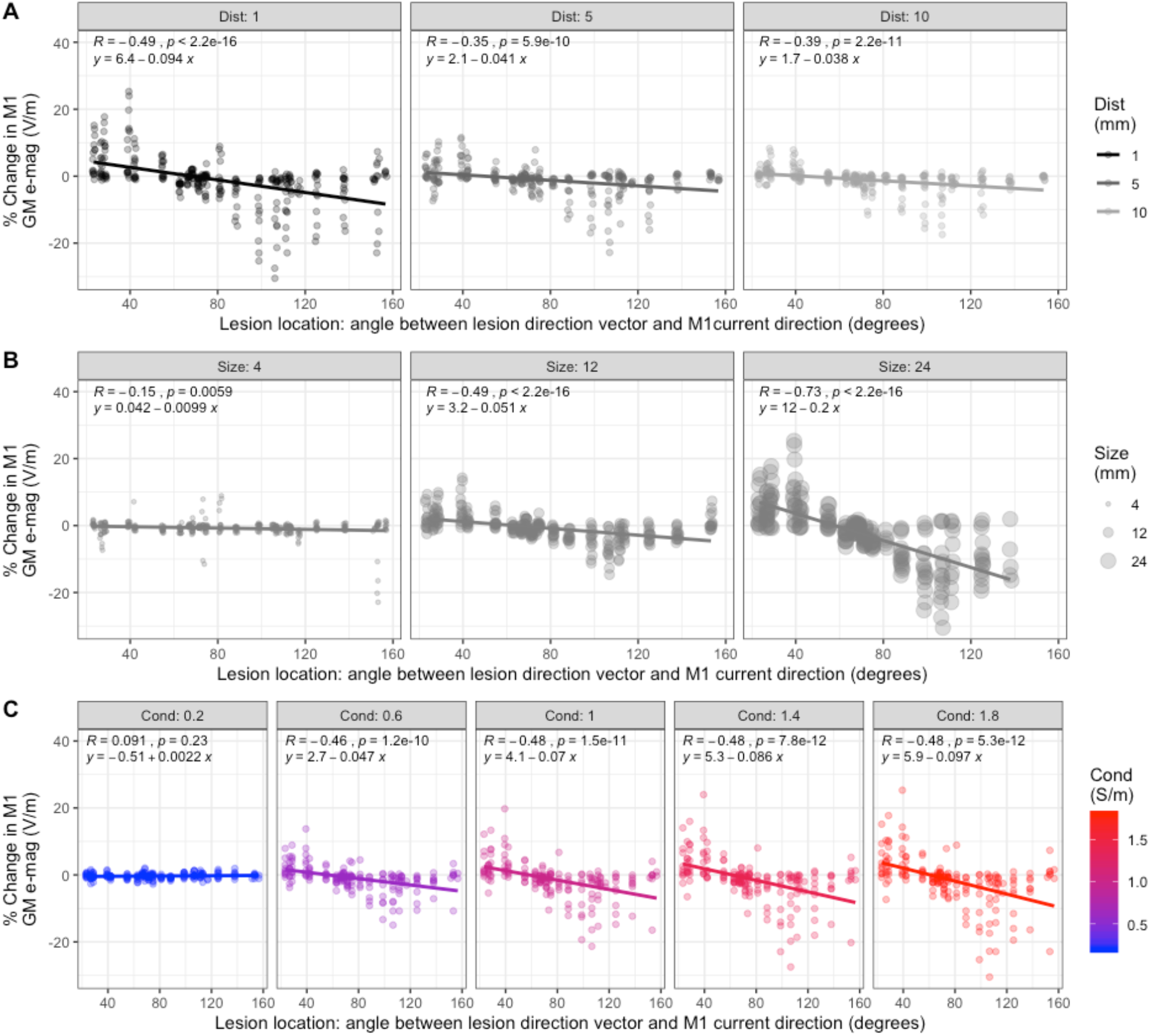
Interaction between the effect of lesion location with distance, size and conductance on change in electric field magnitude (e-mag) in M1. A: Changes in e-mag in M1 grey matter (GM) plotted against lesion location, split by lesion distance (in mm). Data from both participants are displayed, and pooled together for analysis. Lesions located in-line with the predominant orientation of current flow in M1 increased e-mag, whereas those in the opposite direction caused a decrease. This was modulated by lesion distance, where closer lesions to the ROI had a greater impact on e-mag change. B: Changes in e-mag in M1 GM plotted against lesion location, split by lesion size (radius in mm), demonstrating that larger lesions have a greater impact on e-mag change. C: Changes in e-mag plotted against lesion location, split by conductance (in S/m), showing that lesions with higher conductance have a greater effect on e-mag changes.

**Figure 6:**
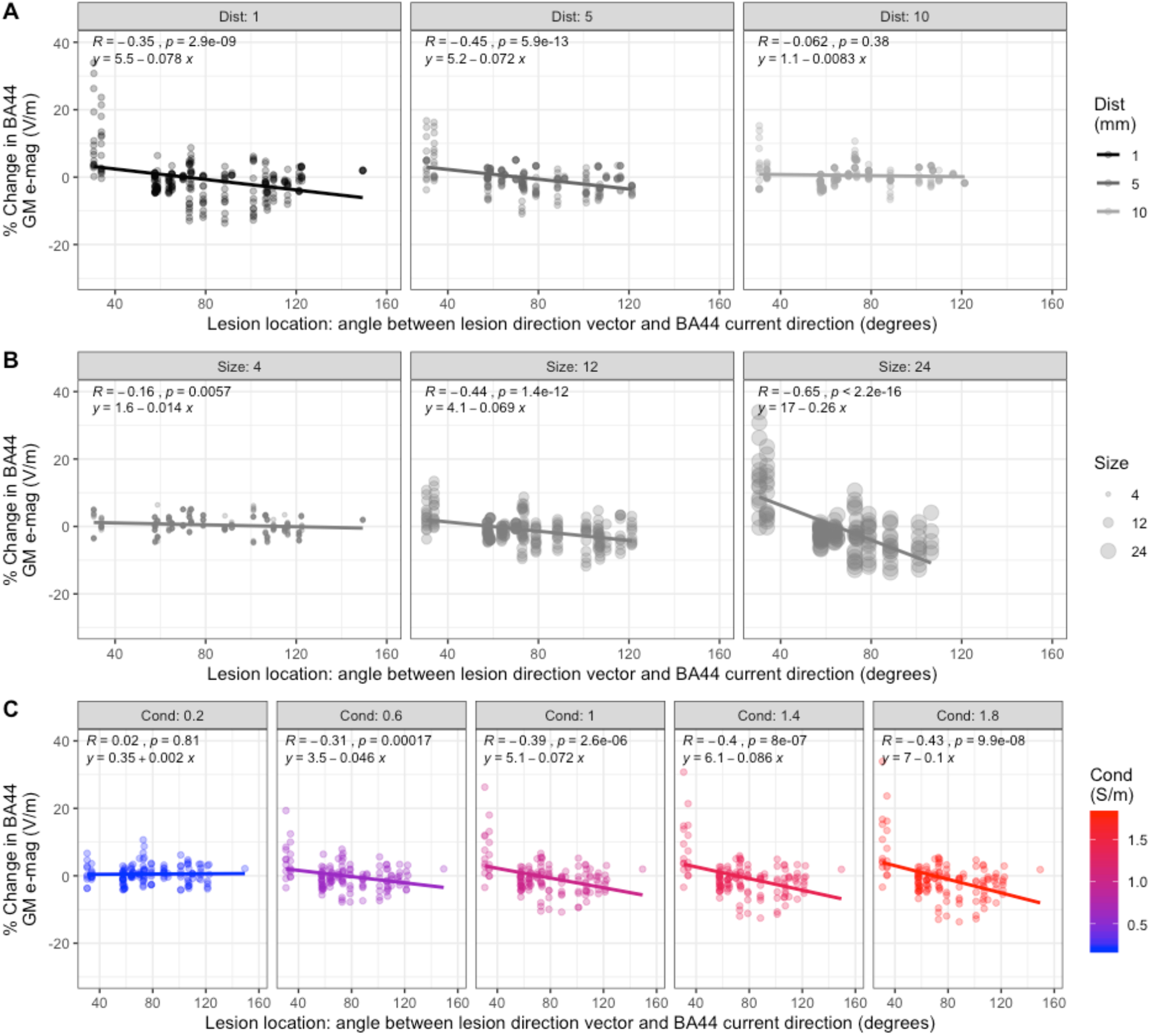
Interaction between the effect of lesion location with distance, size and conductance on change in electric field magnitude (e-mag) in BA44. A: Changes in e-mag in BA44 grey matter (GM) plotted against lesion location, split by lesion distance (in mm). Data from both participants are displayed, and pooled together for analysis. Lesions located in-line with the predominant orientation of current flow in BA44 increased e-mag, whereas those in the opposite direction caused a decrease. This was modulated by lesion distance, where closer lesions to the ROI had a greater impact on e-mag change. B: Changes in e-mag in BA44 GM plotted against lesion location, split by lesion size (radius in mm), demonstrating that larger lesions have a greater impact on e-mag change. C: Changes in e-mag plotted against lesion location, split by conductance (in S/m), showing that lesions with higher conductance have a greater effect on e-mag changes.

Including all the data, from both montages in a single analysis, with montage as a random effect again did not influence the significance of the results. Furthermore, including montage as a fixed effect resulted in no significant main effect of montage and no significant interactions between montage and any of the other effects or 2-way interactions (p>0.25). Once again, this indicates that all effects are consistent across both the M1 and BA44 simulations.

### Optimising tDCS montage in the lesioned brain

To determine whether altering the tDCS montage could compensate for the observed lesion-induced impact on e-mag, we determined the optimal tDCS montages for certain lesion conditions. The target function of ROAST v3.0 was used to identify the optimal montage in the target M1 of participant P01 in the non-lesioned brain as well as with large, close, RAI and LPI lesions (figure 7A). The optimal electrode montage was defined as the pair of electrode locations which maximised electric field magnitude, across all component directions, within the M1 ROI. In the non-lesioned brain, the modelling positioned the anode on CP3 and the cathode on CP4 (CP3-CP4). This montage achieved 0.204 V/m within the target ROI, compared with 0.156 V/m for the conventional M1 montage we initially used (CCP3-Fp2). However, for the lesions used in this step, the optimal electrode locations for maximizing electrical field magnitude were quite different, with CP1-P8 suggested for the RAI lesion condition, and P5-Cz suggested for the LPI lesion. These montages both achieved e-mag of 0.193V/m in the ROI, compared with around 0.182 V/m and 0.129 V/m using the original M1 montage for the equivalent RAI and LPI conditions.

While montage optimisation can compensate for the lesion-induced impact on e-mag, substantially moving the electrodes does result in a change in the direction of current flow through the ROI. Previous studies indicate the importance of the direction of current flow (Bachtiar et al., 2018; Lafon et al., 2017; Rawji et al., 2018). To address this issue we performed roast-target again, this time optimising electrode positions based on maximising only the component of the electric field where current is moving radially into the cortical surface – thought to maximally impact the underlying neurons (Lafon et al., 2017). This resulted in electrode placements that were more similar between lesion conditions (figure 7B). For both the non-lesioned brain and the LPI condition CP1-FT10 was optimal, and for the RAI condition CP1-F10 was optimal, moving the cathode only slightly. Using this more constrained optimisation the electric field magnitude, however, did result in a greater variation in e-mag across the conditions (non-lesioned 0.152V/m; RAI 0.192V/m; LPI 0.124V/m). With all other factors held stable, the electric field magnitude should scale linearly with the applied stimulator current. This means small adjustments to the stimulator output could be used to match the e-mag between conditions (Evans et al., 2020) (see figure 7C).

**Figure 6:**
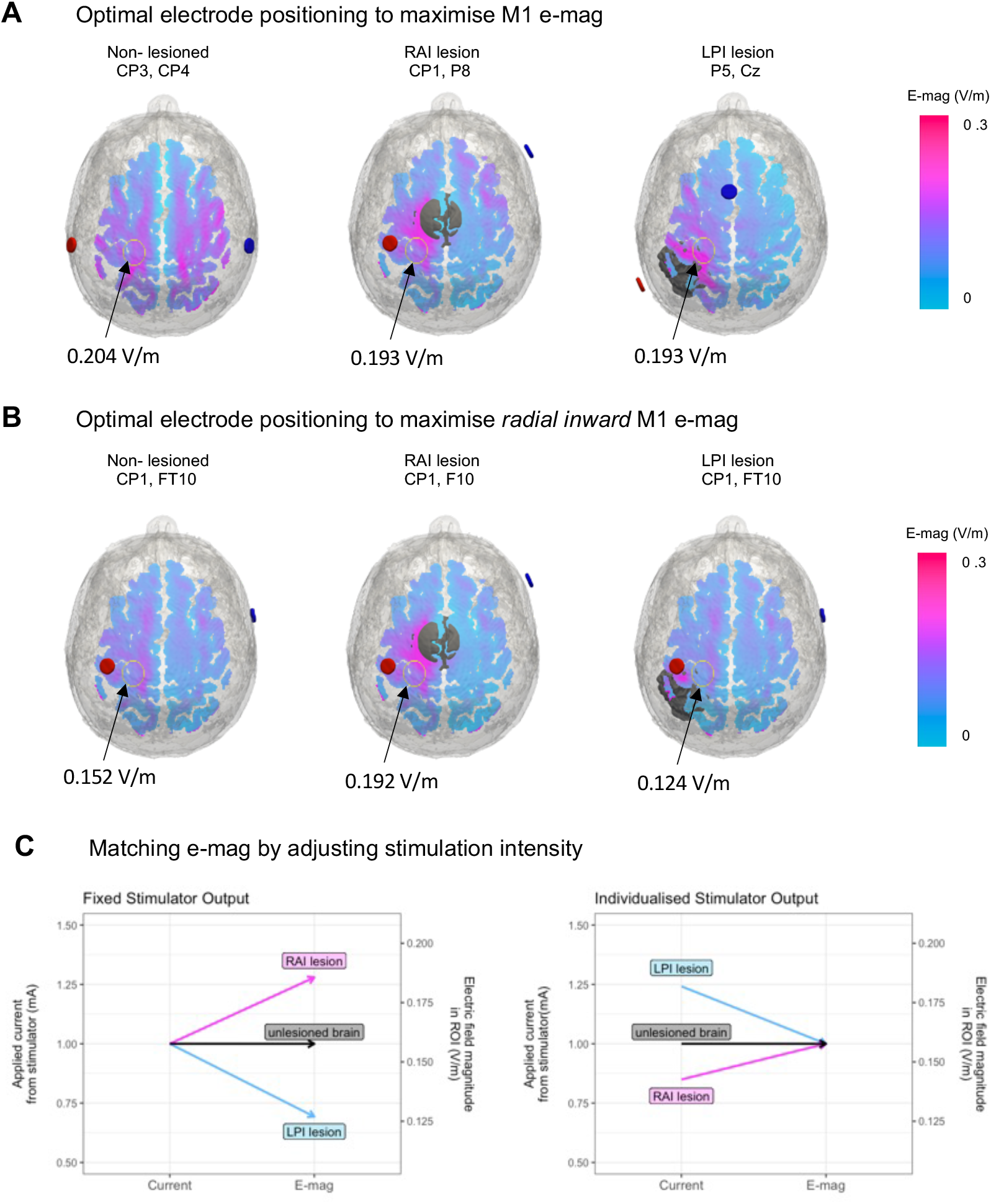
Individualising the tDCS montage and stimulation parameters to match electric field magnitude (e-mag) within M1. A: Optimal electrode locations to maximise the total e-mag in the grey-matter (GM) of the M1 ROI of the non-lesioned brain of participant P01, as well as in the brains with large (24mm radius), close (1mm distance) lesions due right-anterior-inferior (RAI) or left-posterior-inferior (LPI). Lesions were assigned a conductivity of 1.65S/m and tDCS was applied at 1mA. B: Optimal electrode locations to maximise radial inwards flowing e-mag C: Demonstrating how the applied current from stimulator would need to be adjusted in order to achieve the same e-mag within the ROI for each of the displayed lesion locations. Using formula from Evans et al., 2020: *individualised dose = (target e-mag/actual e-mag) x fixed Dose*

## Discussion

In this study, we modelled the effect of synthetic lesions on tDCS-induced e-mag. We found that distance from the ROI, size and conductance of the lesions all impacted the absolute change in e-mag, with increases or decreases of more than 30%. To further probe why some lesions caused increases in e-mag and others decreases, we investigated the effect of lesion location. We found a systematic effect of location, whereby lesions positioned in-line with the predominant orientation of current flow in the ROI, where the ROI was located in between the anode and the lesion, tended to increase the magnitude of current flow within that ROI. Lesions that were positioned maximally out-of-line with the current flow caused a decrease in the magnitude of current flow within the ROI. This effect was modulated by the lesion distance, size and conductance. Lesions that were larger, closer to the ROI, and had a higher conductance tended to have the greatest impact - whether positive or negative - on e-mag. These effects were consistent when examining both a montage targeting M1 and a montage targeting BA44. Additionally, the same pattern of results was seen in both participants (see Supplementary Figures 1 and 2). Taken together, lesion characteristics strongly influence current flow in an individual, an effect that exacerbates inter-individual variability in the delivery of electrical current in stroke. However, our results also point towards a generalisable pattern in the influence of lesions on tDCS-induced e-mag within ROIs which could be used to guide future studies.

The results presented here build on the findings of previous modelling studies examining the effect of lesions on tDCS-induced current flow patterns (Datta et al., 2011; Dmochowski et al., 2013; Galletta et al., 2015; Minjoli et al., 2017; Piastra et al., 2021). These studies used MRI scans of stroke patients and tended to find that lesions caused a decrease in current flow in the region of interest (Minjoli et al., 2017; Piastra et al., 2021). Our results replicate the recent findings by Piastra and colleagues, that larger lesions with higher assigned conductivity have the greatest influence on electric field magnitude (Piastra et al., 2021). Here we also present the novel finding that lesion location, with respect to the direction of current flow, is a key driver in determining the influence of the lesion on current flow, and lesions can cause increases in electric field magnitude in ROIs, as well as decreases.

By modelling synthetic lesions in healthy brains we were able to independently manipulate several lesion parameters to determine their influence on current flow relative to the ‘healthy’ non-lesioned condition and to each other. This enabled control for inter-individual anatomy, which itself affects current flow, and allowed us to use a wide variety of lesion characteristics which could otherwise only be assessed using a high number of real patient scans.

### Implications for clinical applications

A major issue facing the adoption of brain stimulation techniques into clinical practice is the large inter-individual variability in responses (Hordacre et al., 2021). This variability is likely to be driven, at least in part, by differences in current flow through the regions of interest (Laakso et al., 2018). In addition to the high degree of variability seen across healthy individuals (Evans et al., 2020; Laakso et al., 2015), here we show that large lesions will further increase this variability by up to 30%. In the present case, the synthetic nature of our lesions allowed for systematic manipulation of lesion characteristics, but the variability observed here is likely to be amplified in stroke populations with their inherently even more variable lesion characteristics.

Using current flow models to individualise tDCS protocols has been suggested as a method to reduce inter-individual variability (Bestmann and Ward, 2017) either by altering the intensity of stimulation (Evans et al., 2020) or by altering electrode placement (Dmochowski et al., 2011). We used an intuitive and freely available software to perform this individualised optimisation of electrode montage for a non-lesioned brain and the two lesion conditions causing the largest increases or decreases to current flow in the M1 ROI (Dmochowski et al., 2013; Evans et al., 2020; Huang et al., 2019). We found that optimising for electrode position allows for compensating for lesion-induced variability. However, the resulting montages differed largely across conditions. This is not insignificant, as different electrode positions are likely to result in different current directions, albeit similar current strength, in the ROI. Previous studies indicate the importance of the direction of current flow (Bachtiar et al., 2018; Lafon et al., 2017; Rawji et al., 2018). To enable comparable current direction and current strength across conditions, the optimisation was constrained to maximise current flowing radially inward through the cortex - thought to maximally impact the underlying neurons (Lafon et al., 2017). In this case, the optimal electrode configuration was similar across lesion conditions, though the lesions did still cause deviations of around 20% in electric field magnitude. However, as demonstrated by Evans and colleagues, stimulator output can be adjusted in order to match the electric field magnitude in the ROI (Evans et al., 2020). Even for the most extreme conditions here, the maximum applied stimulator current was well within established tDCS safety guidelines (Rossi et al., 2009), making this approach a viable strategy. Taken together, lesion-induced variability can be compensated for by optimising electrode location and stimulator output in each individual based on their specific lesion characteristics.

The optimisation procedures used here are relatively time-consuming and rely on having a high-resolution whole head MRI of each patient post-stroke. While obtaining these scans is possible for some research projects, it may not be feasible for large-scale clinical practice given the availability and cost of high-field MRI. The results presented here could be used, in combination with a 2D MR image, to provide some indication of whether a lesion is likely to impact the electric field magnitude in a region of interest over-and-above the normal inter-individual variability. For example, lesions that are small, distant or lie orthogonal to the path of current flow will have little impact on the electric field magnitude within the region of interest.

### Directions for future research

Our results also highlight the effect of assumptions about the lesion conductance on the degree of electric field change caused by a lesion. In order for current flow simulations to provide accurate estimations of electric field magnitude, it is essential that appropriate conductance values be assigned to lesions. Most previous work has modelled lesions as CSF, which is assigned a very high conductance (Datta et al., 2011; Dmochowski et al., 2013; Galletta et al., 2015; Minjoli et al., 2017). In reality, lesions are not filled with pure CSF and the real conductance is therefore likely to be lower. To the authors knowledge, only one study has investigated the conductance of lesioned tissue using magnetic resonance electrical impedance tomography (MREIT) in a single patient. Here, the predicted conductance value was 1.2 S/m in the lesioned tissue (van Lier et al., 2012). While further research is needed to generate a robust measurement, this value is well below the 1.65 S/m typically assigned to CSF. Our results indicate that lesions with lower conductivity values have less effect, but conductivities in the region of 1.2 S/m can still result in e-mag changes of 20-25%, compared to a non-lesioned brain.

An additional complication is that the area typically segmented as a lesion is not homogenous tissue, but rather has a gradient from maximally damaged pure lesioned tissue, through partially affected perilesional tissue, to healthy tissue (Rekik et al., 2012). Rather than having a constant conductance, it is likely that conductance varies across the lesion. Furthermore, conductance may not be stable across time following the stroke; diffusion MRI metrics, which are known to correlate with conductivity (Tuch et al., 2001, 1999), have been shown to change in the perilesional tissue from acute to chronic stages following stroke (Thiel et al., 2004; van der Zijden et al., 2008). To ascertain the true effect of lesions on tDCS-induced current flow, further work is needed to identify accurate conductivity measures and determine whether these values change across time post stroke, location and distance from the lesion centre.

## Conclusions

In this study, we systematically modelled the influence of synthetic lesions with different locations, sizes and conductivities on the tDCS-induced electric field within two ROIs, for two individuals. A general pattern emerged across both ROIs and individuals. Lesions that were lying in-line with the predominant direction of current flow within the non-lesioned brain tended to increase the electric field magnitude in the ROI, while lesions that were in the opposite direction tended to decrease the electric field in the ROI. This effect was significantly modulated by lesion distance, size and conductance. Lesions that were closer to the ROI, larger in size and had higher conductance induced the greatest e-mag changes, with increases/decreases of up to 30%. These results demonstrate the potential for increased variability in current flow in lesioned brains, which could in turn impact the behavioural effects of tDCS in these patient populations. We also show that small lesions, far from the region of interest, have relatively little impact on electric field magnitude in the ROI. Our results underline the need for improved estimates of conductivity in lesioned tissue if we are to accurately estimate the true path of current flow through the brains of patients and optimise tDCS for clinical practice. Finally, we provide examples of how to individualise tDCS procedures in an effort to maximise current flow through the ROI and reduce variability across individuals.

## Acknowledgements

A Johnstone is funded by the Dunhill Medical Trust [RPGF1810\93]; C Zich [201718-13], C Evans and J Lee [201617-03] are funded by Brain Research UK.

## Supplementary Figures

**Supplementary Figure 1:**
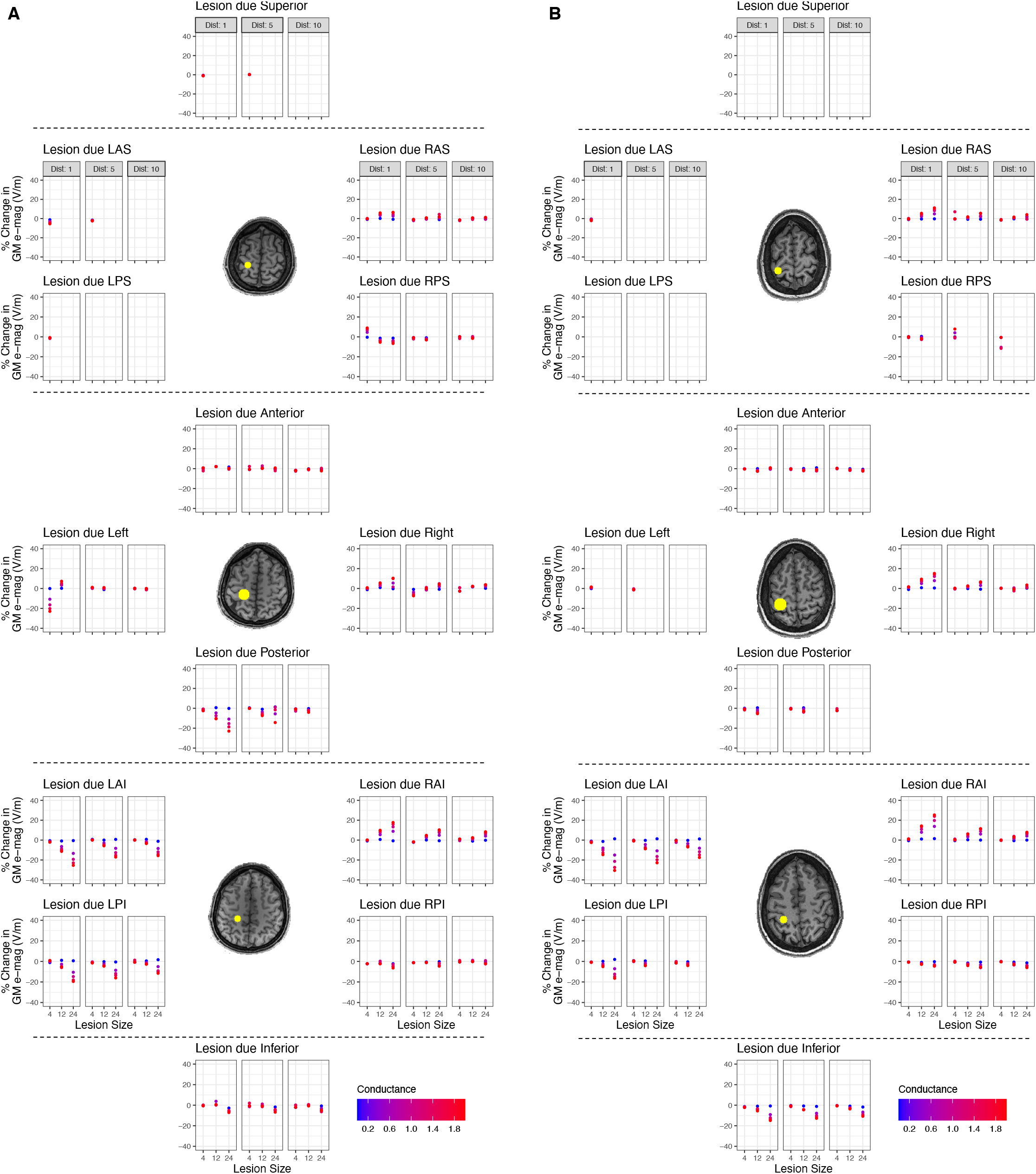
Percentage change in M1 electric field magnitude (e-mag) for all lesions. A: Percentage change in e-mag for all lesion locations, sizes (in mm), distances (in mm), and conductivities (in S/m) for participant P01. B: Same for participant P02

**Supplementary Figure 2:**
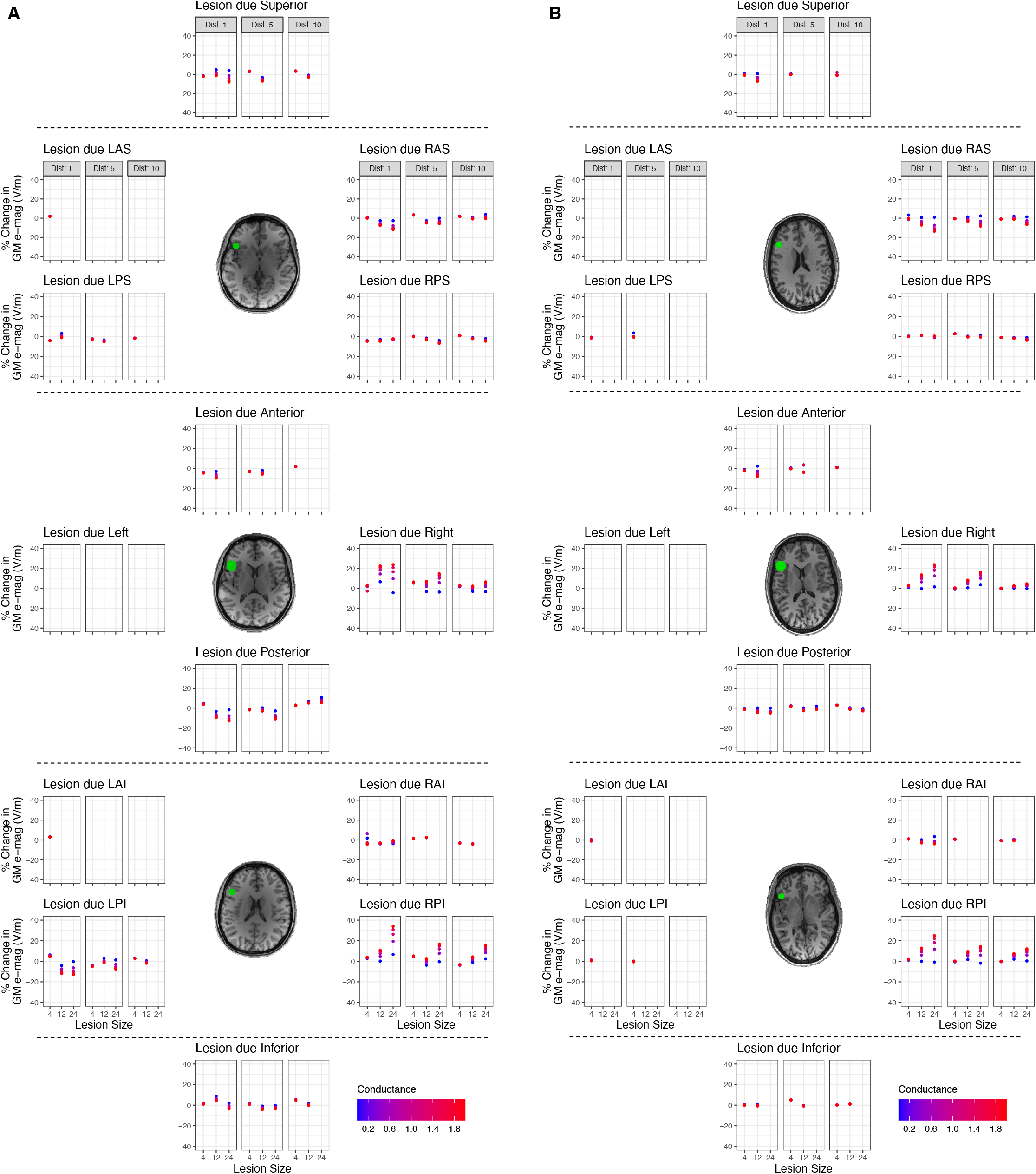
Percentage change in BA44 electric field magnitude (e-mag) for all lesions. A: Percentage change in e-mag for all lesion locations, sizes (in mm), distances (in mm), and conductivities (in S/m) for participant P01. B: Same for participant P02

## Figure Legends

***Figure 1: Modelled lesion parameters***

A: Rendered brain of example participant (P01) with M1 ROI (yellow) and BA44 ROI (green). B: tDCS montage targeting M1, with anode in red, cathode in blue.

C: BA44 tDCS montage.

D: The 14 potential lesion directions N.B. lesions where the central voxel was outside the brain were omitted.

E: The three different lesion distances (1mm, 5mm, 10mm), where distance was measured as the shortest Euclidean distance from the edge of the ROI to the edge of the lesion.

F: The three different lesion sizes, defined by lesion radius (4mm, 12mm, 24mm).

G: The five different lesion conductivities (0.2S/m, 0.6S/m, 1S/m, 1.4S/m, 1.8S/m)

***Figure 2: Current flow modelling***

A: Electric field magnitude (e-mag) image for M1 montage for the non-lesioned ‘healthy’ brain of a single subject. M1 outlined in yellow.

B: E-mag image for same montage but in a lesioned brain (filled white circle; RAI lesion, distance 1mm, size 12mm, conductivity 1.2S/m).

C: Difference image (lesioned minus non-lesioned brain). Note the strong increases and decreases around the lesion, reaching magnitudes of approximately +/-0.1V/m.

D: Electric field direction across the brain, coloured by predominant direction of current flow.

E: Calculation of angle between the lesion direction (black line) and the average direction of electric field within the grey matter of the ROI (orange line).

***Figure 3: Larger lesions, closer to the ROI, with high conductivity have a greater impact on electric field magnitude (e-mag) change within the ROI***

A-C: Scatter plots with mean and standard error (SE) of the *absolute* percentage e-mag changes in M1 caused by lesions with different sizes, distances and conductance. Data are the results from individual simulations and pooled from both participants for analysis.

D-F: Scatter plots with mean and SE of *absolute* e-mag change in BA44 GM. Individual data points are jittered on the x-axis for display purposes.

***Figure 4: Lesion location determined the sign of electric field magnitude (e-mag) change***

A, B: Percentage change in e-mag in M1 region of interest (ROI) caused by lesions due left-anterior-inferior (LAI) or right-anterior-inferior (RAI), with varying distances, sizes and conductivities for participant P01. Data points show mean percentage change across all ROI voxels, error bars show the 16^th^ and 84^th^ percentile values for change across voxels. Inset: Absolute change in e-mag across the whole brain for LAI and RAI lesions of size 12mm, distance 1mm and conductance 1.4S/m.

C,D: Same as above, but for participant P02.

***Figure 5: Interaction between the effect of lesion location with distance, size and conductance on change in electric field magnitude (e-mag) in M1***

A: Changes in e-mag in M1 grey matter (GM) plotted against lesion location, split by lesion distance (in mm). Data from both participants are displayed and pooled together for analysis. Lesions located in-line with the predominant orientation of current flow in M1 increased e-mag, whereas those in the opposite direction caused a decrease. This was modulated by lesion distance, where closer lesions to the ROI had a greater impact on e-mag change.

B: Changes in e-mag in M1 GM plotted against lesion location, split by lesion size (radius in mm), demonstrating that larger lesions have a greater impact on e-mag change.

C: Changes in e-mag plotted against lesion location, split by conductance (in S/m), showing that lesions with higher conductance have a greater effect on e-mag changes.

***Figure 6: Interaction between the effect of lesion location with distance, size and conductance on change in electric field magnitude (e-mag) in BA44***

A: Changes in e-mag in BA44 grey matter (GM) plotted against lesion location, split by lesion distance (in mm). Data from both participants are displayed and pooled together for analysis. Lesions located in-line with the predominant orientation of current flow in BA44 increased e-mag, whereas those in the opposite direction caused a decrease. This was modulated by lesion distance, where closer lesions to the ROI had a greater impact on e-mag change.

B: Changes in e-mag in BA44 GM plotted against lesion location, split by lesion size (radius in mm), demonstrating that larger lesions have a greater impact on e-mag change.

C: Changes in e-mag plotted against lesion location, split by conductance (in S/m), showing that lesions with higher conductance have a greater effect on e-mag changes.

